# Determining Optimal Placement of Copy Number Aberration Impacted Single Nucleotide Variants in a Tumor Progression History

**DOI:** 10.1101/2024.03.10.584318

**Authors:** Chih Hao Wu, Suraj Joshi, Welles Robinson, Paul F. Robbins, Russell Schwartz, S. Cenk Sahinalp, Salem Malikić

**Author notes:** Joint first authors (Chih Hao Wu, Suraj Joshi). Joint last authors (S. Cenk Sahinalp, Salem Malikić).

## Abstract

Intratumoral heterogeneity arises as a result of genetically distinct subclones emerging during tumor progression. These subclones are characterized by various types of somatic genomic aberrations, with single nucleotide variants (SNVs) and copy number aberrations (CNAs) being the most prominent. While single-cell sequencing provides powerful data for studying tumor progression, most existing and newly generated sequencing datasets are obtained through conventional bulk sequencing. Most of the available methods for studying tumor progression from multi-sample bulk sequencing data are either based on the use of SNVs from genomic loci not impacted by CNAs or designed to handle a small number of SNVs via enumerating their possible copy number trees. In this paper, we introduce `DETOPT`, a combinatorial optimization method for accurate tumor progression tree inference that places SNVs impacted by CNAs on trees of tumor progression with minimal distortion on their variant allele frequencies observed across available samples of a tumor. We show that on simulated data `DETOPT` provides more accurate tree placement of SNVs impacted by CNAs than the available alternatives. When applied to a set of multi-sample bulk exome-sequenced tumor metastases from a treatment-refractory, triple-positive metastatic breast cancer, `DETOPT` reports biologically plausible trees of tumor progression, identifying the tree placement of copy number state gains and losses impacting SNVs, including those in clinically significant genes.

## A Introduction

Cancer progression is a product of an evolutionary process through which the concomitant accumulation and selection of genomic aberrations in somatic cells promote their survival and unregulated proliferation [23]. Tumors typically develop into genetically heterogeneous subpopulations of cells (subclones), and those with more diverse intratumoral compositions may have an increased likelihood of acquiring treatment resistance and metastatic potential. Reconstructing the evolutionary progression history of heterogeneous tumors not only could inform risk stratification for targeted therapies, but may also reveal mutational mechanisms of resistance, as these would correspond to the positive selection and continual evolution of particular subclones over others [19].

While single-cell DNA sequencing has advanced studies of tumor progression substantially [22], most newly generated DNA sequencing datasets are still obtained through conventional and more cost-effective bulk DNA sequencing. In particular, multiple bulk-sequenced samples obtained from one or more tumor biopsies of the same cancer patient provide robust signals for studying a tumor’s progression history and can help in identifying subclonal mutations conferring treatment resistance and/or driving metastatic spread. From a spatial point of view, sampling across multiple regions of the same tumor allows for the profiling of subclones occurring in distinct tumor regions that would not be recoverable in a single sample [9]. Likewise, longitudinal sampling across multiple timepoints can be informative about how tumors change in composition from a temporal perspective [31,1]. The clonal composition of a tumor may change (e.g., in response to a therapy) from an earlier time point with distinct dominant subclones, to later timepoints with emerging subclones dominating the tumor [31].

The existing computational methods for inferring tumor progression trees (also known as trees of tumor evolution) aim to integrate multiple bulk-sequenced samples in a number of ways. However, most of these methods such as PhyloSub [11], LiCHeE [25], CITUP[17], AncesTree [6], PASTRI [29], CALDER [21], MIPUP [10] and Pairtree [34] are primarily designed for somatic single nucleotide variants (SNVs) located in genomic loci not impacted by copy number aberrations (CNAs). Although there have been some developments in the design of computational methods that account for SNVs from copy number altered mutational loci (such as SPRUCE [7]), these methods either operate on a small set of variants and/or (sub)clones [7,8] or make restrictive assumptions about the possible relationships between SNVs and copy number gains and losses [5]. More recently, PACTION [28], a method for integration of trees independently derived by the use of CNAs (CNA-tree) and SNVs (SNV-tree) was proposed. While there is a history of methods for bulk sequencing data that focus specifically on CNA-trees [32,12], inference of CNA-trees is still a largely unsolved problem. In the original study [28] the authors performed exhaustive search over all possible CNA-trees, which limits practical applications of this method to tumors characterized by low numbers of CNA clones. DeCiFer [30] is another recent method, which clusters SNVs based on their read counts while accounting for the impact of CNAs. While descendant cell fractions of SNVs reported by this method can in principle be used as the input to several of the available tree inference methods (e.g., CITUP or AncesTree), DeCiFer does not impose a joint tree of tumor progression shared by all variants, which can result in decreased accuracy of the inferred trees.

There are several challenging problems encountered in the computational integration of SNVs and CNAs for tree inference. One particular limiting factor to the development of methods capable of handling CNA-impacted SNVs was the lack of scalable approaches for detailed copy number profiling of tumors with multiple bulk-sequenced samples. One notable recent development in this direction is HATCHet [35] which successfully leverages complementary signals from multiple bulk samples to provide a comprehensive subclonal architecture of a tumor with respect to CNAs.

In this work we introduce a combinatorial optimization method, `DETOPT`, for determining optimal placement in tumor progression history of SNVs from the genomic regions impacted by CNAs. `DETOPT` utilizes disjoint genomic segments (herein, identified by HATCHet), each with a consistent CNA profile across all subclones of the tumor, to simultaneously place CNAs and SNVs from regions impacted by CNAs onto a tree of tumor progression inferred by the use of SNVs from diploid regions. Specifically, the placement of variants reported by `DETOPT` aims to minimize both the difference between the observed and inferred variant allele frequencies of the CNA-impacted SNVs and difference between the input and inferred allelic fractional copy numbers of CNA-impacted segments in the tumor across all samples.

On simulated multi-sample bulk sequencing data, we demonstrate that `DETOPT` outperforms available alternatives achieving a better accuracy in capturing lineage relationships among SNVs. We also present `DETOPT`’s results on previously unpublished multi-region longitudinal bulk whole-exome sequencing data obtained from multiple tumor metastases in a patient which was not only refractory to previous lines of endocrine therapy and immunotherapy, but whose cancer subsequently progressed after partial response to adoptive cell therapy. `DETOPT` inferred that several clinically relevant neoantigenic and treatment resistance-mediating mutations have been subject to allele-specific CNAs, corroborating the hypothesis that SNVs can work in tandem with allele-specific CNAs to direct tumor progression. `DETOPT` is based on a time-efficient Integer Linear Programming formulation and is freely available at https://github.com/algo-cancer/DETOPT.

## 2 Methods

### 2.1 Tree representation of tumor progression

In this work, we represent the progression history of a tumor as a rooted tree *T* (Figure 1a)^1^, with a set of nodes *V* (*T*), where the root node *r* represents a (sub)population of normal cells and each of the remaining nodes represents a subclone, i.e. a genetically uniform (sub)population of tumor cells that are distinct from other subclones. Because each distinct subclone corresponds to a distinct node of *T* and vice versa, considering the normal cells as one subclone, in the remainder of this work we use the terms subclone and node interchangeably. We also assume that the root node is free of somatic genomic aberrations, whereas every other node *v* is associated with at least one distinct somatic CNA or SNV that is represented in *T* on the edge connecting *v* to its parent *p*(*v*). These somatic genomic aberrations (including both CNAs and SNVs) distinguish subclone *v* from its parental subclone *p*(*v*); the set of somatic aberrations harbored by *v* can be obtained by combining somatic aberrations of *p*(*v*) and the set of somatic aberrations represented on the edge connecting *v* to *p*(*v*) (below we also refer to the somatic aberrations represented on this edge as aberrations that occur for the first time at node/subclone *v*).

**Fig. 1:**
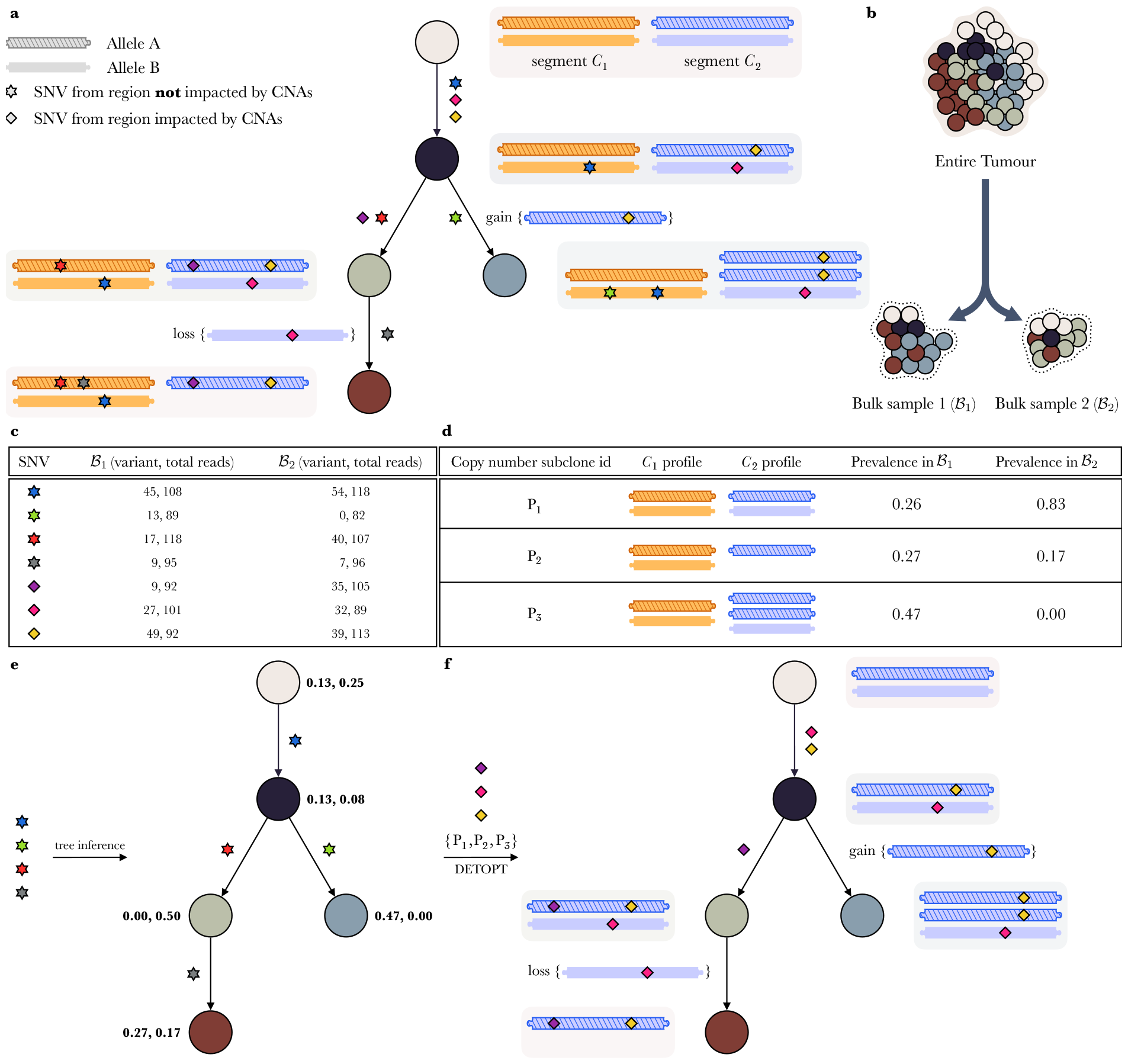
Overview of tumor progression history and its reconstruction using our method `DETOPT`. **a**. A tree of tumor progression. Each node corresponds to a genetically distinct population of cells (subclone). SNVs from regions not impacted by CNAs are represented as stars and SNVs from regions impacted by CNAs as diamonds, all of distinct colors. The root node represents a population of healthy cells that is free of somatic SNVs and CNAs and any other node (subclone) differs from its parent by the set of SNVs and CNAs present on the edge connecting them. In this figure we show only two genomic segments, one that is not impacted by CNAs (orange) and another that undergoes copy number gains and losses (blue). Next to each node, copy number profile of the corresponding subclone is shown in a shaded rounded rectangle. **b**. An example of the input to a bulk sequencing experiment consisting of two tumor samples obtained from different regions of the same tumor biopsy. (Note that, in addition to sampling different regions of the same tumor, multiple tumor samples can also be obtained by taking biopsies at different timepoints or from different anatomic sites, e.g., primary and metastasis.) **c**. Read count data obtained for each SNV in each bulk sample. The first and second numbers in each pair respectively represent the number of reads supporting the variant and total coverage at the mutated site in the corresponding sample. **d**. Subclonal architecture of the sequenced tumor samples inferred with respect to CNAs. Note that three subclones in the tree shown in panel (a) share the same copy number profile implying that there exist three copy number subclones (i.e., genetically distinct subpopulations of cells with respect to copy number profiles). For each of the copy number subclones, its proportion in each of the samples is inferred. **e**. A tree of tumor progression inferred using one of the well established methods designed for SNVs from genomic regions not impacted by CNAs. The first and the second number shown next to each node represent cellular prevalence of the corresponding subclone in samples *ℬ*_1_ and *ℬ*_2_, respectively. **f**. Placement in the tree of tumor progression of copy number gains and losses of genomic segments and SNVs that they harbor using `DETOPT`. In this panel we only show the placements that are inferred by `DETOPT`. Taking the union of trees in panels (e) and (f) yields complete tumor progression history on the entire set of SNVs and CNAs, together with cellular prevalence values of all subclones across both sequenced samples.

### 2.2 Input data and preprocessing

We assume that we are given bulk DNA sequencing data of *h* tumor samples from the same cancer patient, denoted *ℬ* = {*ℬ*_1_, *ℬ*_2_, …, *ℬ*_*h*_}, obtained by targeted, whole-exome or whole-genome sequencing (Figure 1b).

We preprocess the given sequencing data first by using some existing mutation caller, to obtain a set *M* = {*M*_1_, *M*_2_, …, *M*_*m*_} of somatic SNVs, each reported to be present in at least one of the samples *ℬ*_1_, *ℬ*_2_, …, *ℬ*_*h*_. For each SNV *M*_*i*_ and each sample *ℬ*_*s*_, we also calculate a number of variant reads supporting *M*_*i*_, denoted as *v*_*is*_, as well as the total number of reads spanning the genomic position of *M*_*i*_, denoted as *t*_*is*_ (Figure 1c).

For the next preprocessing step we use some of the sophisticated tools (e.g. recently developed HATCHet [35]) to infer subclonal architecture of samples *ℬ*_1_, *ℬ*_2_, …, *ℬ*_*h*_ with respect to copy number states (Figure 1d). Specifically, we use these tools to partition the genome into non-overlapping segments *C* = {*C*_1_, *C*_2_, …, *C*_*q*_}, where each segment corresponds to contiguous genomic loci with the same number of “A” and “B” alleles in any cell. Here, for any given segment *C*_*j*_, alleles A and B refer to the two copies of this segment on a pair of autosomal chromosomes in normal diploid cells. For any segment *C*_*j*_, a normal cell typically contains one copy of allele A and one copy of allele B, while among the tumor cells the number of copies of these alleles might vary due to CNAs. We denote by *𝒮* (*C*_*j*_) the set of pairs of numbers (*i*_*A*_, *i*_*B*_), respectively corresponding to the number of alleles A and B of segment *C*_*j*_, that exist in a given tumor as reported by a tool for subclonal architecture inference with respect to copy number states. For example, in the example shown in Figure 1, we have *𝒮* (*C*_2_) = {(1, 1), (1, 0), (2, 1)} implying that in the sequenced tumor samples, with respect to segment *C*_2_, there exist three distinct cell (sub)populations consisting of: (i) cells with 1 copy of allele A and 1 copy of allele B (ii) cells with 1 copy of allele A and 0 copies of allele B (iii) cells with 2 copies of allele A and 1 copy of allele B. In the remainder of the paper we call these “CNA-consistent” segments of the tumor.

After completing the above steps, the set of SNVs can be divided into sets *M*′ and *M* − *M*′, where *M*′ consists of all SNVs that belong to genomic segments *C*′ ⊆ *C* that are not impacted by CNAs. For a given SNV *M*_*i*_ ∈ *M*, let *V AF* (*M*_*i*_, *ℬ*_*s*_) = *v*_*is*_*/t*_*is*_ denote its observed variant allele frequency in sample *ℬ*_*s*_. If the genomic locus of *M*_*i*_ is diploid and free of CNAs in all tumor cells (i.e., *M*_*i*_ ∈ *M*′), assuming that *M*_*i*_ is heterozygous^2^, the fraction of cells in *ℬ*_*s*_ that harbor *M*_*i*_, commonly referred to as *observed cellular prevalence* of *M*_*i*_, can be estimated as 2 *· V AF* (*M*_*i*_, *ℬ*_*s*_). A similar expression can easily be derived for SNVs from haploid regions of the genome (e.g. X and Y chromosomes in males) assuming that they are free of CNAs. Existing methods for inferring progression history of a tumor by the use of SNVs from genomic loci not impacted by CNAs [11,25,6,17,21] typically aim to come up with tree *T* where (i) each edge is associated with at least one distinct SNV from the set *M*′, and (ii) each node *v* is associated with *h*-dimensional vector *ϕ*_*v*_ = [*ϕ*_*v*1_, *ϕ*_*v*2_, …, *ϕ*_*vh*_]^*T*^, where *ϕ*_*vs*_ represents the inferred prevalence of node/subclone *v* in sample *ℬ*_*s*_. To solve this problem, they employ different optimization techniques, usually with a goal of finding a solution that minimizes total or limits maximum absolute difference between the observed and inferred cellular prevalence values of the SNVs. One example of such tree is shown in Figure 1e.

### 2.3 Placement of CNA-impacted SNVs in tumor progression history using the available methods

Determining placement in tumor progression history of SNVs from genomic regions impacted by CNAs is a challenging task as there is usually no straightforward relationship between the observed read counts and (unknown) cellular prevalence values of SNVs from such genomic regions [4,33]. CNAs overlapping with a given SNV can impact its read counts in multiple distinct ways depending on the relative placement of the SNV and CNAs in the tumor progression history, as well as whether the allele to which SNV belongs is gained/lost during a copy number event [5]. For such SNVs, existing tree reconstruction tools typically rely on clustering methods that can estimate cellular prevalence values by *correcting* the observed read counts for CNAs. Two examples of such methods are PyClone [27] and the more recent and more general DeCiFer [30], which can leverage information on tumor subclonal architecture inferred by tools like HATCHet to provide better corrections, especially for deletion events. However, neither of these methods infers a tumor progression tree; rather, they cluster variants and correct for CNAs without imposing a joint tree structure on all variants. On the other hand, the group of methods that integrate SNVs and CNAs in inferring a progression tree typically focus on a small number of variants or clones [7,8] or assume restrictive set of possible relationships between SNVs and CNAs [5].

### 2.4 Description of DETOPT

In DETOPT, we assume that we are given the following as input: (i) read counts for SNVs from regions impacted by CNAs (ii) subclonal architecture of the sequenced samples with respect to CNAs, and (iii) a tree of tumor progression *T*, together with node/subclone frequencies *ϕ*, inferred on the set *M*′ of SNVs from genomic regions not impacted by CNAs (see Section 2.2 and Figure 1c-e). We also assume that tree *T* captures all subclones present in the sequenced tumor samples (see Section 4 for a discussion of this assumption).

For each CNA-consistent segment *C*_*j*_ from the set *C* − *C*′ (recall that *C* − *C*′ represents the set of all CNA-consistent segments impacted by CNAs), let *ℳ* (*C*_*j*_) denote the set of all SNVs located in *C*_*j*_. Given the above described input, for each *C*_*j*_ ∈ *C* − *C*′, the goal of `DETOPT` is to: (i) assign a copy number state from set *𝒮* (*C*_*j*_) to each node of *T*, (ii) assign each SNV *M*_*i*_ located in *ℳ* (*C*_*j*_) to one edge (equivalently, vertex) in *T*, and (iii) assign each SNV *M*_*i*_ ∈ *ℳ* (*C*_*j*_) to one of the alleles A and B of *C*_*j*_, together with, for each node of *T*, the number of its copies that harbor *M*_*i*_ at that node (see Figure 1f). For a given segment *C*_*j*_ ∈ *C* − *C*′, `DETOPT`’s objective is to minimize the weighted sum across all samples of: (i) absolute difference between the observed and inferred variant allele frequencies across all SNVs from *ℳ* (*C*_*j*_), and (ii) absolute difference between the input total fractional copy numbers of segment *C*_*j*_ and those implied by `DETOPT`-assignment of copy number states from *𝒮* (*C*_*j*_) to nodes of tree *T*. For each segment we formulate this as an instance of Integer Linear Programming (ILP) and solve it by the use of an ILP solver (in this work we used Gurobi). For simplicity of notation and without loss of generality, below we focus on an arbitrary segment *C*_*j*_ ∈ *C* − *C*′ and assume that *ℳ* (*C*_*j*_) = {*M*_1_, *M*_2_, …, *M*_*l*_}. In addition, whenever we refer to allele A or B, we are referring to the allelic copies of segment *C*_*j*_.

First, for each SNV *M*_*i*_ ∈ *ℳ* (*C*_*j*_) and node *v* ∈ *V* (*T*), we introduce a binary variable *δ*_*iv*_, which is set to 1 if and only if *M*_*i*_ occurs for the first time at node *v* (more precisely on the edge connecting *p*(*v*) and *v*). As the root *r* of *T* represents the (sub)population of normal cells that are assumed to not contain somatic aberrations, we set *δ*_*ir*_ = 0 for all *i*. Similarly as discussed above, we also assume that the genomic locus of *M*_*i*_ is “hit by” a mutation exactly once (although, as will be seen below, we allow existence of multiple copies of the variant allele harboring *M*_*i*_ acquired through gains of *C*_*j*_). This can be modeled with the following constraint

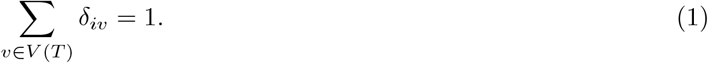

Similarly, for each copy number state *S*_*k*_ ∈ *𝒮* (*C*_*j*_) and each node *v* ∈ *V* (*T*) we introduce a binary variable *θ*_*kv*_ that is set to 1 if and only if *S*_*k*_ occurs for the first time at node *v*. Thus we add the constraint

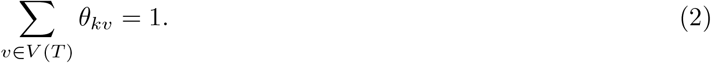

To ensure that each node *v* has a unique copy number status we add the following constraint^3^

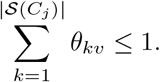

Assuming (without loss of generality) that *S*_1_ denotes the copy number status of segment *C*_*j*_ in the normal cells, we require that *θ*_1*r*_ = 1.

Next, for each *v* ∈ *V* (*T*) and each *k* ∈ {1, 2, …, |*𝒮* (*C*_*j*_)|} let *ψ*_*vk*_ denote a binary variable that indicates copy number status at node *v*. More precisely, *ψ*_*vk*_ is set to 1 if and only if copy number status at node *v* is *S*_*k*_. The values of variables *ψ* are entirely defined by the structure of a given tree *T* and the assignment of variables *θ* as follows: first, if copy number state *S*_*k*_ occurs for the first time at *v* (i.e., *θ*_*kv*_ = 1) then we require that *ψ*_*vk*_ = 1, which can be ensured by adding constraint

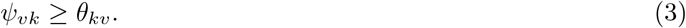

If *v* is a non-root node such that no copy number state from *𝒮* (*C*_*j*_) occurs for the first time at *v* and copy number status at *p*(*v*) is *S*_*k*_, then the copy number status at *v* must also be *S*_*k*_, which can be ensured by constraint

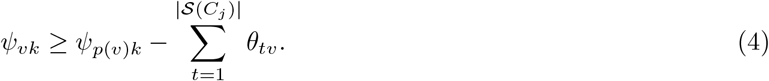

It can be easily verified that, for any given *v*, the above constraints on *ψ* would ensure that one of the variables *ψ*_*vk*_ is set to 1. To ensure that the copy number status at *v* is unique we add constraint

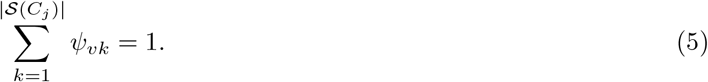

For each node *v*, let *𝒯*_*Av*_ denote a non-negative integer variable encoding the total number of copies of allele *A* at node *v*. Then the value of *𝒯*_*Av*_ is given by the formula

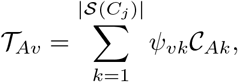

where *𝒞*_*Ak*_ denotes the total number of copies of allele *A* in the copy number state *S*_*k*_ (note that the value of *𝒞*_*Ak*_ is known since it is a part of the input to `DETOPT`). We define variables *𝒯*_*Bv*_ analogously.

When assigning copy number states to tree nodes, if at some node the number of copies of allele *A* reaches zero then the number of copies of allele *A* in all of its descendants must also be zero. This can be ensured by adding constraint

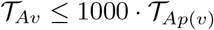

for each non-root node *v*. Analogous constraints are added for *B* allele. Here, 1000 represents a large constant that exceeds the maximum number of copies of allele *A* or *B* so that the above constraint is trivially satisfied if *𝒯*_*Ap*(*v*)_ *>* 0. It will also be used in several constraints below, whereas its multiplicative inverse 0.001 will be used to represent a small constant. While these constants are used solely to make it easier to follow the presented constraints, in practice, for a given *C*_*j*_, 1000 can be replaced by any number greater than *c* and 0.001 by any positive number less than 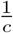, where *c* represents the maximum allelic copy number of *C*_*j*_ among all subclonal populations and can be obtained from *𝒮* (*C*_*j*_).

Next, for a given variant *M*_*i*_ ∈ *ℳ* (*C*_*j*_), let *µ*_*iA*_ denote the binary variable which is set to 1 if and only if *M*_*i*_ occurs for the first time on allele A. Analogously we define *µ*_*iB*_. In order to ensure that *M*_*i*_ is present only on copies of allele A or only on copies of allele B, we add the following constraint

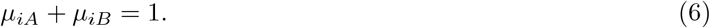

Now, for each node *v* ∈ *V* (*T*) we introduce non-negative integer variables *A*_*iv*_ and *B*_*iv*_ which respectively represent the number of copies of alleles A and B harboring *M*_*i*_. As the root node *r* represents the normal cells, which do not harbor somatic aberrations, we set *A*_*ir*_ = *B*_*ir*_ = 0. To ensure that the number of the variant copies of allele does not exceed its total number of copies, and that the number of the variant copies of allele not harboring *M*_*i*_ is set to zero, we add the following constraints^4^

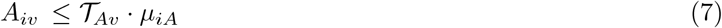

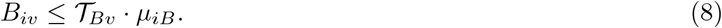

Next, we require that there is at least one variant copy of allele harboring *M*_*i*_ at the node of its occurrence. Thus we add the following constraints

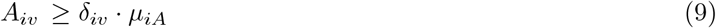

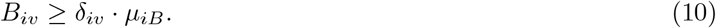

For a given non-root node *v*, when comparing the numbers of copies of allele *A* at *v* and *p*(*v*) these numbers can be equal, or the number of copies at *v* is smaller (i.e., there is copy number loss of one or multiple copies of allele *A* at *v*) or greater (i.e., there is copy number gain of one or multiple copies of allele *A* at *v*). Since these three cases have different implications for modeling some of the constraints given below, we introduce variables *e*_*Av*_, *l*_*Av*_ and *g*_*Av*_ that encode which of these three cases we have at *v*. More precisely, *e*_*Av*_ is set to 1 if and only if we have equal number of copies of allele *A* at *v* and *p*(*v*), *l*_*Av*_ is set to 1 if and only if we have loss and *g*_*Av*_ is set to 1 if and only if we have gain of copies of this allele at *v*. Note that values of *e*_*Av*_, *l*_*Av*_ and *g*_*Av*_ depend on other unknown variables of our model and constraints that ensure that they take desired values are provided in Supplementary Section A.1.

Now, consider a non-root node *v* and its parent *p*(*v*). Focusing on allele A, we have three possible cases at node *v* compared to *p*(*v*):

#### Case 1

Copy number status of allele A is equal at *v* and *p*(*v*), that is *e*_*Av*_ = 1.

In this case the number of copies of A harboring *M*_*i*_ at *v* is equal to that at *p*(*v*), except if *M*_*i*_ occurs for the first time at *v*. To enforce these we add the following constraints

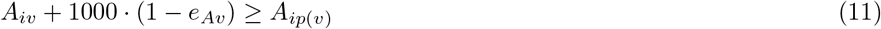

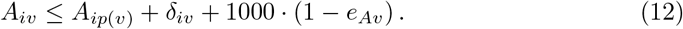

Note that the terms 1000 · (1 − *e*_*Av*_) are added in order to make both of the above constraints trivially satisfied when we are not in this case (i.e., if *e*_*Av*_ = 0).

#### Case 2

There is a loss of one or multiple copies of allele A occurring between *p*(*v*) and *v*, that is *l*_*Av*_ = 1.

To ensure that in this case the total number of variant copies of allele A harboring *M*_*i*_ does not increase compared to the parent node, except if *M*_*i*_ occurs for the first time at *v*, we add the following constraint

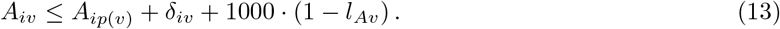

#### Case 3

There is a gain of one or multiple copies of allele A occurring between *p*(*v*) and *v*, that is *g*_*Av*_ = 1.

First, to ensure that in this case the number of copies of allele *A* harboring *M*_*i*_ at *v* is not lower than at *p*(*v*), we add constraint

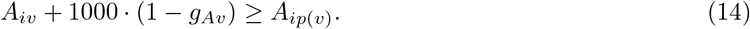

Next, we add constraint

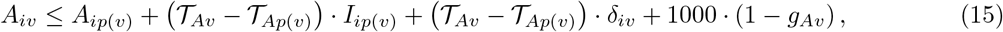

where *I*_*ip*(*v*)_ denotes *indicator* variable that is set to 1 if and only if *M*_*i*_ is present at node *p*(*v*), or, in other words, if there is at least one copy of allele *A* or allele *B* harboring *M*_*i*_ at *v* (constraints ensuring that variables *I*_*iv*_ take desired values are provided in Supplementary Section A.2). Note that the above constraint ensures that, if the variant is present at *p*(*v*) (i.e., *I*_*ip*(*v*)_ = 1, which also implies *δ*_*iv*_ = 0) then the total number of copies of *A* harboring *M*_*i*_ at *v* is no greater than the total number of such copies at *p*(*v*) plus the number of copies of *A* that were gained. On the other hand, if there are no copies of *A* harboring *M*_*i*_ at *p*(*v*), then such copies can be present at *v* only if *M*_*i*_ occurs for the first time at *v* (i.e., if *δ*_*iv*_ = 1, which implies *A*_*ip*(*v*)_ = 0 and *I*_*ip*(*v*)_ = 0) and we allow to have at most *𝒯*_*Av*_ − *𝒯*_*Ap*(*v*)_ of them, with the exact number depending on the relative timing of *M*_*i*_ and copy number gains.

One important subcase to consider in the case of a gain is the following: assume that there is a gain of allele A at some node *v* and that all copies of A at *p*(*v*) harbor *M*_*i*_. In this case we require that all copies of A at *v* also harbor *M*_*i*_. To ensure this, we first introduce a set of binary variables *U*_*Aiv*_ for each variant *M*_*i*_ and each node *v*, with *U*_*Aiv*_ = 1 if and only if at node *v* there is a non-zero number of copies of allele A and all of them harbor *M*_*i*_ (i.e., *U*_*Aiv*_ = 1 if and only if *𝒯*_*Av*_ = *A*_*iv*_ *>* 0). Constraints ensuring that *U*_*Aiv*_ take desired values are provided in Supplementary Section A.3. Now, in addition to constraints (14) and (15) added above, we also add the following constraint

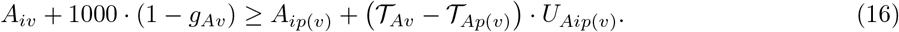

Analogous set of variables and constraints are added for allele B.

Finally, our objective is to minimize the (weighted) sum of: (i) difference between the observed and tree-implied variant allele frequencies, summed across all variants *M*_1_, *M*_2_, …, *M*_*l*_ and samples *ℬ*_1_, *ℬ*_2_, …, *ℬ*_*h*_, and (ii) difference between the observed and input total fractional allelic copy numbers of segment *C*_*j*_ across all samples *ℬ*_1_, *ℬ*_2_, …, *ℬ*_*h*_. Below, the input total fractional allelic copy numbers of alleles *A* and *B* of segment *C*_*j*_ in sample *ℬ*_*s*_ are denoted as *ℱ*_*A*_ (*C*_*j*_, *ℬ*_*s*_) and *ℱ*_*B*_ (*C*_*j*_, *ℬ*_*s*_), respectively, and we assume that they are obtained from the output of a copy number caller (e.g., for the example shown in Figure 1d we have *ℱ*_*A*_ (*C*_2_, *ℬ*_1_) = 0.26 · 1 + 0.27 · 1 + 0.47 *·* 2 = 1.47). More formally, our objective is minimizing

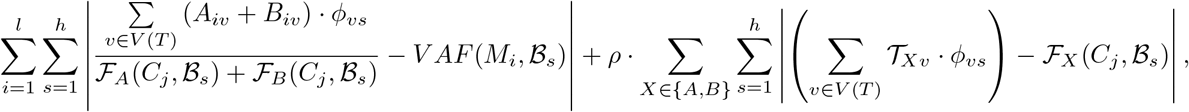

where *ρ* as a regularization parameter introduced to balance the two objective terms, which we estimate by assessing `DETOPT`’s performance over multiple values of *ρ* on simulated data. Generally, we find that setting *ρ* to 0.25 optimizes `DETOPT`’s performance on simulations.

The above procedure is repeated for all segments *C*_*j*_ to obtain a complete tree of tumor progression involving all SNVs and segmental copy number changes.

## 3 Results

### 3.1 Benchmarking on Simulated Data

We first assessed the performance of `DETOPT` on a simulated data. In order to compare its performance against the available alternatives, in addition to running `DETOPT`, we also ran PhyloWGS [5], as well as combination of DeCiFer [30] and CITUP [17]. PhyloWGS is a tree inference method which is primarily based on the use of read count data derived from SNVs, but can also account for the impact that CNAs can have on the read counts of the SNVs. DeCiFer is more recently developed clustering method that groups SNVs based on their descendant cell fractions, which can serve as an input for tree inference method CITUP. To the best of our knowledge, these are currently the best available alternatives that can be used for understanding placement in tumor progression history of SNVs impacted by CNAs. Our comparisons are based on two commonly used measures of accuracy:

1. *Ancestor-Descendant Accuracy* of CNA-impacted SNVs defined as the proportion of SNV pairs with at least one being CNA-impacted, which have an ancestor-descendant relationship in the ground truth tree that is preserved in the inferred tree (after the assignment of CNA-impacted SNVs to nodes).
2. *Different-Lineage Accuracy* of CNA-impacted SNVs defined as the proportion of SNV pairs with at least one being CNA-impacted, which are in different lineages in the ground truth tree, that are also in different lineages in the inferred tree (after the assignment of CNA-impacted SNVs to nodes).

Using the previously described simulator [16], we generated data with varying number of bulk samples *h* ∈ {3, 5, 7, 10}. In order to simulate tumors with different rates of copy number instability, we also varied the fraction of genome impacted by CNAs between 0.1, 0.2 and 0.4. For simplicity, below we will refer to this fraction as CNA rate. In the results shown in Figure 2, all simulated trees were of size 10, with total number of mutations *m* = 500 and depth of sequencing coverage equal to 100*×*. As can be seen from this figure, `DETOPT` consistently outperforms the combination of DeCiFer and CITUP, as well as PhyloWGS, in terms of both measures. Notably, while the performance of `DETOPT` remains stable across varying CNA rates, we observe that performance of DeCiFer+CITUP deteriorates with increased CNA rates. As expected, both `DETOPT` and DeCiFer+CITUP show improved performance with increase in the number of simulated samples. On the other hand, PhyloWGS in most cases shows improved different-lineage accuracy with increased CNA rates, however this comes at a cost of decreased ancestor-descendant accuracy suggesting that it tends to report more branching tree topologies as CNA rates increase. Surprisingly, in a substantial number of instances the performance of PhyloWGS deteriorates when number of samples is increased from 7 to 10.

**Fig. 2:**
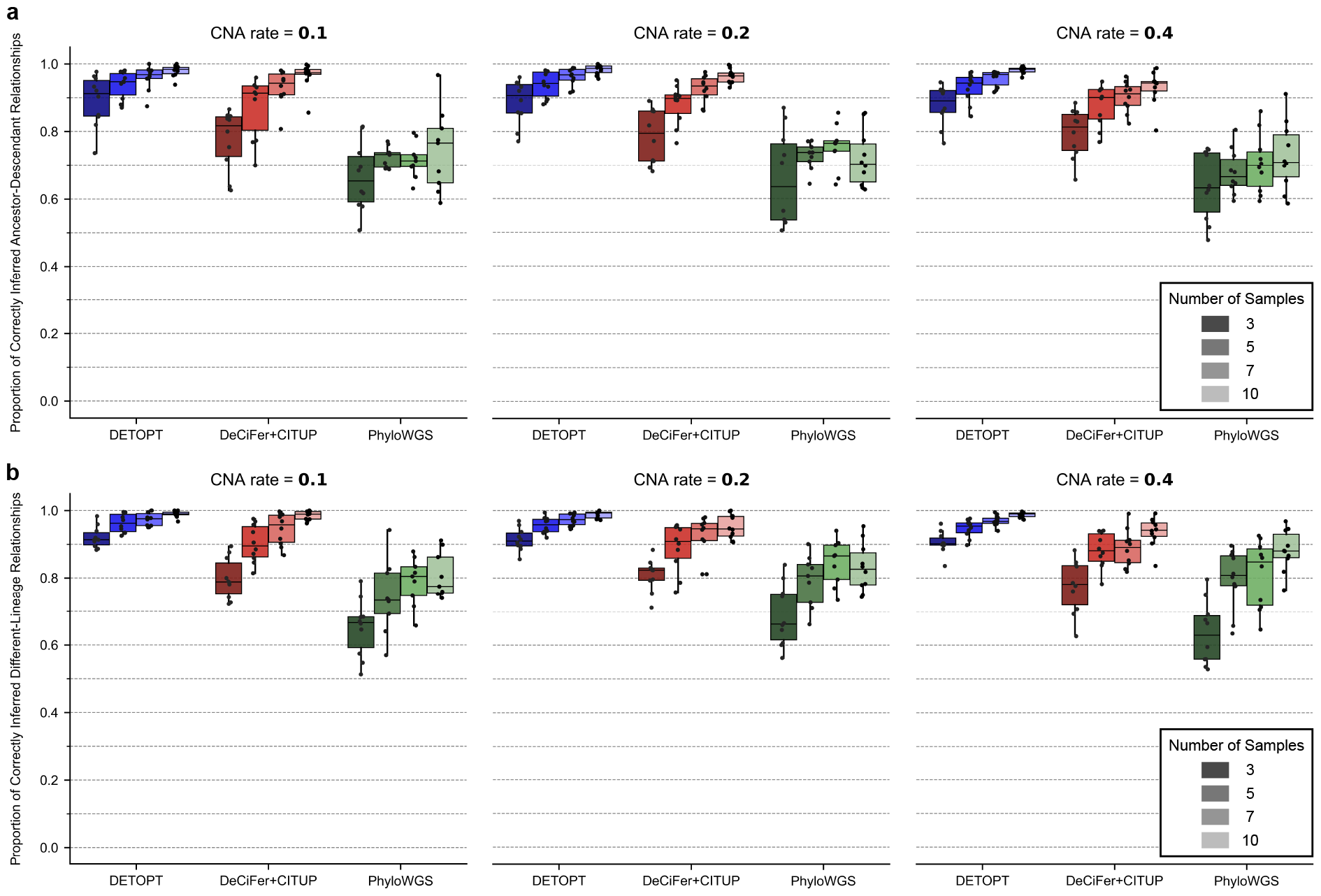
Comparison of `DETOPT`, DeCiFer+CITUP, and PhyloWGS on simulated data. CNA rate denotes the fraction of SNVs impacted by CNAs. In all simulated instances shown in this figure, the total number of SNVs was set to 500, tree size to 10, and sequencing coverage to 100*×*. For each parameter combination, 10 different trees were simulated. Plotted are the **a**. Ancestor-Descendant accuracy (see the main text for definition). **b**. Different-Lineage accuracy (see the main text for definition) for each tool. PhyloWGS, which uses Markov Chain Monte Carlo (MCMC) sampling to cluster mutations and infer the tree, failed to converge in 5 (out of 120) instances within the time limit of 120 hours and no results for this method are shown for those 5 instances.

In addition to varying number of bulk samples and CNA rates, we also assessed robustness of all methods to varying tree sizes and number of mutations. Due to space constraints, results of these additional comparisons are shown in Supplementary Figures 1-4.

### 3.2 Application to Real Data

We analyzed an ER+/PR+/Her2+ metastatic breast cancer (mBrCa) tumor refractory to previous passive immunotherapies, hormone- and targeted-therapies (from patient 4355 from [36]). This patient relapsed 10 months after partial response on adoptive cell therapy (ACT) using autologous tumor-infiltrating lymphocytes (TILs) concomitantly with anti-PD-1 immunotherapy following nonmyeloablative lymphodepleting chemotherapy. The progression of this tumor is thought to have been mediated by allele-specific copy number gains and loss of heterozygosity (LOH) events.

In a previously unpublished multi-sample bulk whole exome sequencing dataset, we had 18 tumor fragments that were sampled from recurring and new metastatic lesions in lymph node, liver, lungs, and breast at four different time points corresponding to pre- and post-TIL-ACT (−212, +81, +342, and +523 days relative to therapy) and all were bulk whole-exome sequenced.

We first built the baseline tree on 325 copy number neutral SNVs as described in Supplementary Section C.1. We observe that the inferred tree features very strong concordance between the observed and inferred cellular prevalence values of mutational clusters (see Supplementary Figure 5). Moreover, we emphasize that this base tree also has longitudinal consistency. The inferred structure of subclones is concordant with the time points from which samples (containing these subclones) were collected, even though we have not included time of sampling as a constraint during our tree inference. In this base tree, we observe that subclones from later samples only emerge from subclones present at earlier time points and the subclones from samples at the latest sampling time point appear towards the leaves of the tree (see Supplementary Section C.3).

Given the baseline tree, `DETOPT` was used to infer the placement of 139 SNVs impacted by CNAs, including the neoantigens that were identified in the prior work [36]. In Figure 3, we focus on the placements of SNVs in 8 genes that are identified to have putative role(s) in tumor progression (*PIK3CA, CBFA2T3, JAK2, ANKRD11*) [37,20], acquired treatment resistance (*ESR1*) [24], or tumor immunogenicity (*ANO1, TRIM3, RCL1*) [36]. Note that details about accessing a complete list with placement of all 139 SNVs, as well as the placement of SNVs used for building the baseline tree, are provided in Supplementary Section C.3.

**Fig. 3:**
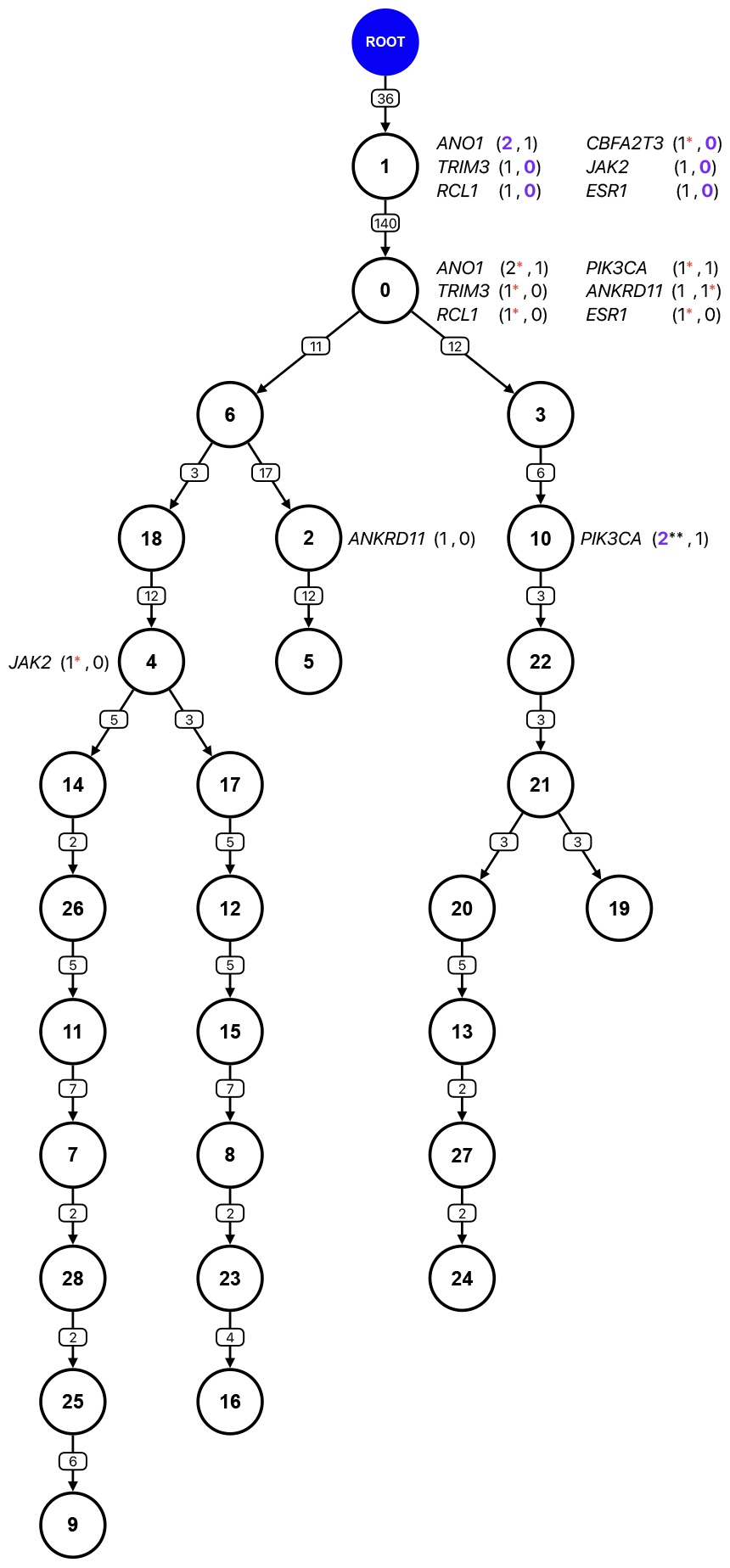
Tumor progression tree with placements of SNVs (from regions impacted by CNAs) inferred by `DETOPT` from multi-region longitudinal bulk whole-exome sequencing data for a patient with triple-positive mBrCa (patient 4355 in [36]). On each edge, the number of copy number neutral mutations are shown. Each node is labeled with a subclone id (note that subclone ids are not given in any particular order). The copy number state of a gene is given as *GENE NAME* (copies of allele A, copies of allele B). Copy number gain and loss events are indicated in purple and occur on the allele that is colored. Asterisks indicate the subclone in which an SNV was acquired (and on which allele the mutation is present on); no asterisk is shown if the mutation was lost due to LOH event e.g., *ANKRD11*. Additional asterisks denote a gain of the mutant allele e.g., *PIK3CA* (2**, 1) indicates gain of mutated allele resulting in two mutant copies harboring mutation in gene *PIK3CA*. Red asterisks indicate the subclone in which the mutation first appeared. For each gene, the root node has 1 copy of allele *A* and 1 copy of allele *B*.

Among the neoantigens identified as therapeutic targets in [36], we found that three fall into segments impacted by CNAs (*ANO1, TRIM3*, and *RCL1*) while the remaining were from diploid regions (*LMB2, INF2, PIGG, SPAG1*). The authors in [36] additionally found that each of these neoantigens were present in all time points in every sampled metastatic lesion, even in those emerging after therapy, suggesting that: (i) these neoantigenic mutations are likely clonal, and (ii) if a loss-of-heterozygosity event occurred, the mutant allele was not impacted. As can be seen from Figure 3, `DETOPT` placed the mutational events for these CNA-impacted neoantigens in the trunk of the tumor progression tree (for all three, and subsequent to the CNA event) and also inferring that the mutations on *TRIM3* and *RCL1* were present on the single remaining allele, consistent with the presence of these mutations across all time points. Further, we compared these placements to those of the copy number neutral neoantigens (*LMB2, INF2, PIGG, SPAG1*) which were placed independently and had the same presence in all samples as the CNA-impacted neoantigens. The placement of these copy number neutral mutations as also clonal/truncal mutations corroborates that `DETOPT` accurately identified the neoantigenic mutations at CNA-impacted sites as clonal/truncal events.

From here, we further extend the results from the prior work, first focusing on the base tree to suggest additional insights into the evolution of these tumor metastases. In this branched tree, we observed that a subtree rooted at subclone 6 (including subclones 2, 5, 18, 4) was present in a metastasis sampled at +81 days; however, these subclones became completely absent in a new metastatic deposit at +342 days, but then present in another, also new metastatic deposit at +523 days (lost subtree rooted by subclone 2 but expanded into a subtree rooted by subclone 4). This gave an example that was consistent with the perplexingly partial response to neoantigen-reactive TILs as previously reported. Investigating this, we identified that *BTF3* was another copy number neutral neoantigen from the prior study; however, unlike other neoantigens, it was first completely absent from two post-ACT samples (+342 days), and then present again in other samples from different metastases (+523 days). And notably, the *BTF3* mutation was placed into subclone 6 and the presence-absence profile of this neoantigenic mutation completely overlapped that of the subtrees which were pruned, entirely or incompletely, to suggest its probable contribution in the elimination of subclones via ACT-TIL. Intriguingly, these samples were both metastases taken from newly developed subcutaneous breast nodules. Since the subtree rooted by the subclone 6 is present in samples taken at +81 days, it likely seeded both these metastases. This is perhaps indicative of a differential response to therapy between metastatic sites. We speculate that the recency of the metastasis (at +342 days) sensitised this particular tumor metastasis while concurrently, other sites which may be older and/or larger, continued to acquire mutations and copy number alterations, for instance, giving rise to the expanded subtree rooted at subclone 4.

Next, using the inferred placements returned by `DETOPT`, we investigated how (sub)clonal mutational and copy number alteration events in mBrCa might drive cancer progression towards treatment-resistant and/or more aggressive phenotypes.

*ESR1* mutations and copy number gains/losses have, together, been well-characterised as a key mechanism of endocrine resistance in HR+ mBrCAs [24,3]. We inferred that *ESR1* acquired the SNV (L79H) following an LOH, placing both as truncal events; coincidentally, this patient was refractory to aromatase inhibitors (i.e., endocrine therapy). *CBFA2T3* (D366N) has been identified as a putative tumor suppressor gene (TSG) shown to be targeted by a loss of 16q as an early event in breast tumorigenesis [13]. We inferred that this gene was impacted by a truncal mutation event that coincided with an LOH event that deleted the functional copy of the gene, likely exacerbating the effect of the mutation. *PIK3CA* mediates regulation of apoptosis and cell cycle progression and mutations in this gene have been implicated tumor initiation and proliferation and metastasis [26,38]. We inferred that the SNV (H1047L), a gain-of-function hotspot variant, was acquired in subclone 0 as a truncal event; then, in subclone 10, there was a gain of the mutant allele as a subclonal event. *ANKRD11* has been shown to be a putative TSG with its down-regulation associated with breast tumorigenesis [15]. We placed the mutation (S769X) in subclone 0, and interestingly, the mutant allele was lost due to an LOH event in subclone 2. *JAK2* acquired an SNV after treatment in samples from a subcutaneous breast metastasis resected at/after +81 days after ACT-TILs (samples from +81 and +523 days). Mutations in *JAK2* have been suggested to contribute to further tumor progression through disruption of interferon (IFN)-*γ*-mediated anti-tumor immune signalling pathways in a known mechanism of acquired resistance to immunotherapy [2]. We inferred that in subclone 1, there was a truncal LOH event; and then in subclone 4, the SNV (C961F) was as a subclonal event. Additionally, we knew that the mutational event for *JAK2* and LOH event for *ANKRD11* were not only subclonal, but also acquired in subclones from samples corresponding only to later time points. Correspondingly, we observed that `DETOPT` was making inferences in a manner that is consistent with the timing of the samples i.e., the tool did not assign a mutation or CNA to a subclone that was not present in the samples. Specifically, it did not place these post-treatment aberrations in a subclone that emerged prior to those from post-treatment.

Notably, computed over all assignments of copy number aberrant SNVs, the average discrepancy between the inferred and observed VAFs was less than 0.04. Given the inferred tumor progression tree, and previous studies on this patient, our results suggest that DETOPT assigns clinically relevant somatic variants to plausible/expected nodes.

### 3.3 Running time

`DETOPT` is highly time-efficient with respect to placing CNA-impacted SNVs in a given tumor progression tree. Most of the running time of our pipeline is spent on pre-processing steps, including the base tree inference from SNVs not impacted by CNAs. For example, on a standard personal laptop, for each of the problem instances presented in this work, `DETOPT` completes assignment of all CNA-impacted variants within less than one minute.

## 4 Discussion and Future Work

In this work we introduced `DETOPT`, a combinatorial optimization approach for time efficient placement of SNVs from genomic segments impacted by CNAs onto a tree of tumor progression using multi-sample bulk DNA sequencing data. Tumor progression tree topology inference from SNVs that are not impacted by CNAs is a well studied computational problem with several efficient solutions. Our contribution extends the usability of these solutions by enabling the integration of CNA-impacted SNVs to obtain progression history that includes the entire set of SNVs detected in the sequenced samples.

On simulated data, we showed that `DETOPT` provides a more accurate placement of CNA-impacted variants compared to the available alternatives. Additionally, on an in-house collected cancer dataset, `DETOPT` was able to place variants that were impacted by genomic gains and losses in plausible (sub)clonal expansions. These results suggest that `DETOPT` may provide novel insights into the progression and heterogeneity of existing and future multi-sample bulk sequenced tumor data sets.

There are several potential directions for future work. (1) `DETOPT`’s ability to assign allele-specific copy number gains and losses to distinct subclones in a tumor progression tree is highly relevant to tumors whose progression is thought to be mediated by allele-specific LOH events. Several tumors in our in-house breast cancer cohort (not presented in this work) feature both clonal and subclonal allele-specific LOH events, which may have contributed to tumor immune evasion - even though the role of LOH events in the evolution of solid epithelial tumors is relatively unknown. Upon obtaining accurate LOH calls, we plan to use `DETOPT` to assign these events to the previously inferred tumor progression trees. (2) While in its current implementation `DETOPT` provides placements of CNA events on a tree of tumor progression with several biologically motivated constraints, it handles each CNA-consistent segment independently. We anticipate that our approach could benefit from a joint consideration of all segments; this, however, would require more comprehensive models of tumor progression and would require further experimental validation. (3) Another extension to `DETOPT` would be to permit the assignment of a segmental CNA state to subclones from distinct tree lineages. Even though it is straightforward to add this feature to the ILP formulation, identifying segments that acquire the same copy number state at different subclones independently is very challenging and would also come at a cost of increased running time. (4) In its current version, `DETOPT` assumes that the tree of tumor progression inferred on SNVs from regions not impacted by CNAs captures all major subclonal populations present within the sequenced tumor samples. While this assumption is plausible for most whole exome/genome sequencing datasets (where even for the subclones whose emergence is driven by CNAs or CNA-impacted SNVs we still expect the presence of passenger SNVs from regions not impacted by CNAs that distinguish these subclones from their parents), in practice it might be violated for targeted sequencing datasets or for tumors with very low SNV mutational burden. In such cases, `DETOPT` could benefit significantly from extending it to enable expansion of the given input tree by addition of new nodes that are distinguished from their parents and/or children solely by CNAs or CNA-impacted SNVs. (5) An additional direction for future work is the integration of bulk and single-cell sequencing data, which is already performed by several available tools on data comprised of SNVs that are not impacted by CNAs [16], or (a limited number of) SNVs impacted by losses only [18], or CNA events alone [14]. Note that `DETOPT` in its current form is primarily applicable to tumors that are moderately impacted by CNAs; this is because it relies on SNVs that are not impacted by CNAs for infering a base tree topology. For tumors exhibiting extensive CNAs that affect most of the genome, it is advisable to perform single-cell sequencing or use tools specifically designed for inferring phylogenies from CNAs in order to obtain a reliable base tree topology.

## Acknowledgements

This work utilized the computational resources of the NIH Biowulf high-performance computing cluster (http://hpc.nih.gov) and Gurobi (http://www.gurobi.com) to solve optimization problems.

## Supplementary Materials

### A Additional Constraints for the ILP formulation

#### A.1 Constraints related to variables *e*_*Av*_, *l*_*Av*_ and *g*_*Av*_

Recall from Section 2.4 that, for each non-root node *v* (and assuming a given CNA-consistent segment *C*_*j*_), binary variable *e*_*Av*_ is required to be set to 1 if and only if the number of copies of allele *A* is equal at *v* and its parent *p*(*v*). In other words, *e*_*Av*_ = 1 if and only if *𝒯*_*Av*_ = *𝒯*_*Ap*(*v*)_. To ensure that *e*_*Av*_ takes desired value, it is sufficient to add the following constraints

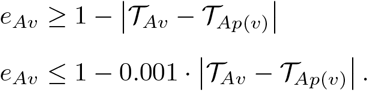

Next, binary variable *l*_*Av*_ is required to be set to 1 if and only if there is a loss of copies of *A* at *v* compared to *p*(*v*) (i.e., *𝒯*_*Av*_ *< 𝒯*_*Ap*(*v*)_). To ensure that *l*_*Av*_ takes desired value, we add the following constraints

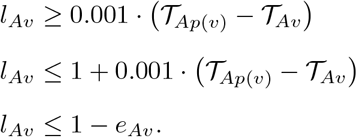

Lastly, binary variable *g*_*Av*_ encodes the remaining, third case, where we have gain of copies of allele *A* at *v* (i.e., *𝒯*_*Av*_ *> 𝒯*_*Ap*(*v*)_). To ensure that this variable takes desired value it is sufficient to add constraint

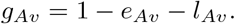

Note that for allele *B* we define analogous variables denoted as *e*_*Bv*_, *l*_*Bv*_ and *g*_*Bv*_.

#### A.2 Constraints related to variables *I*_*iv*_

Recall from Section 2.4 that the *indicator* variable *I*_*iv*_ is a binary variable which encodes whether *M*_*i*_ is present at node *v*. More precisely, *I*_*iv*_ is set to 1 if and only if at node *v* the total number of copies of alleles *A* or *B* harboring *M*_*i*_ is greater than 0. In the absence of deletion events, for a given tree *T, I*_*iv*_ is a linear combination of variables *δ*. However, due to possible presence of deletion events, which can cause losses of all allelic copies harboring *M*_*i*_, we add the following constraints to ensure that *I*_*iv*_ is set to the desired value

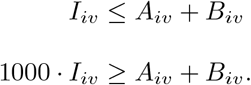

#### A.3 Constraints related to variables *U*_*Aiv*_

Recall from Section 2.4 that *U*_*Aiv*_ is a binary variable that is set to 1 if and only if the number of copies of allele *A* harboring *M*_*i*_ at node *v* is greater than zero and equals the total number of copies of *A* at node *v* (i.e., *𝒯*_*Av*_ = *A*_*iv*_ *>* 0).

We first ensure that *U*_*Aiv*_ is set to 0 if *𝒯*_*Av*_ ≠ *A*_*iv*_. Observe that, due to the way we define variables *𝒯*_*Av*_ and *A*_*iv*_, we have that *𝒯*_*Av*_ ≠ *A*_*iv*_ is equivalent to *𝒯*_*Av*_ − *A*_*iv*_ *>* 0 so in order to ensure that *U*_*Aiv*_ takes desired value in this case it is sufficient to add constraint

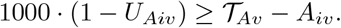

Next, to ensure that *U*_*Aiv*_ is set to 0 in case where no copies of allele *A* at *v* harbor *M*_*i*_ (i.e., *A*_*iv*_ = 0), we add constraint

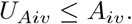

Lastly, if *𝒯*_*Av*_ = *A*_*iv*_ and *A*_*iv*_ *>* 0 then *U*_*Aiv*_ is expected to be set to 1, which can be achieved through constraint

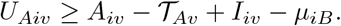

Note that addition of term −*µ*_*iB*_ to the right hand side of the last constraint is necessary due to the possibility that *I*_*iv*_ is set to 1 because *M*_*i*_ is present at *v*, but on allele *B*.

### B Supplementary Tables

Below we present a table containing short description of symbols and variables (labels) used in this work.

**Supplementary Table 1:**
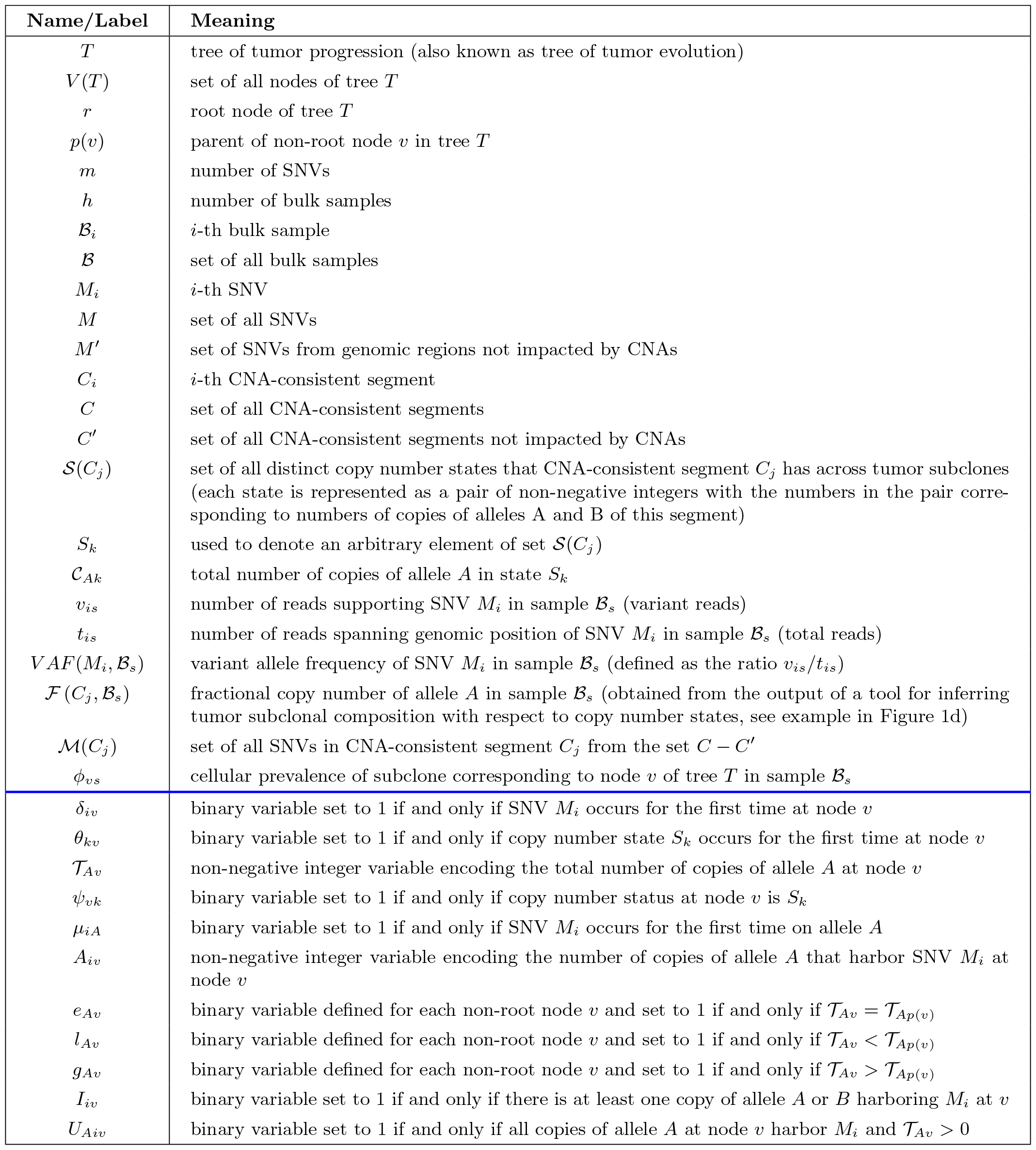
List of variables and labels used in this paper together with their brief descriptions. Note that variables *𝒞*_*Bk*_, *𝒯*_*Bv*_, *µ*_*iB*_, *B*_*iv*_, *e*_*Bv*_, *l*_*Bv*_, *g*_*Bv*_, *U*_*Biv*_ are defined for allele *B* analogously as their matching variables for allele *A*. The values of all variables listed above the blue line are known based on the input to `DETOPT`. All variables below the blue line are unknown and we aim to set their values by solving the ILP presented in Section 2.4. Note also that all variables below the blue line are assumed to be with respect to a given CNA-consistent segment *C*_*j*_ from the set *C* − *C*′ and the set *ℳ* (*C*_*j*_) that consists of SNVs with genomic positions belonging to *C*_*j*_.

| Name/Label | Meaning |
| --- | --- |
| $T$ | tree of tumor progression (also known as tree of tumor evolution) |
| $V(T)$ | set of all nodes of tree $T$ |
| $r$ | root node of tree $T$ |
| $p(v)$ | parent of non-root node $v$ in tree $T$ |
| $m$ | number of SNVs |
| $h$ | number of bulk samples |
| $\mathcal{B}_i$ | $i$-th bulk sample |
| $\mathcal{B}$ | set of all bulk samples |
| $M_i$ | $i$-th SNV |
| $M$ | set of all SNVs |
| $M'$ | set of SNVs from genomic regions not impacted by CNAs |
| $C_i$ | $i$-th CNA-consistent segment |
| $C$ | set of all CNA-consistent segments |
| $C'$ | set of all CNA-consistent segments not impacted by CNAs |
| $\mathcal{S}(C_j)$ | set of all distinct copy number states that CNA-consistent segment $C_j$ has across tumor subclones (each state is represented as a pair of non-negative integers with the numbers in the pair corresponding to numbers of copies of alleles A and B of this segment) |
| $S_k$ | used to denote an arbitrary element of set $\mathcal{S}(C_j)$ |
| $\mathcal{C}_{Ak}$ | total number of copies of allele A in state $S_k$ |
| $v_{is}$ | number of reads supporting SNV $M_i$ in sample $\mathcal{B}_s$ (variant reads) |
| $t_{is}$ | number of reads spanning genomic position of SNV $M_i$ in sample $\mathcal{B}_s$ (total reads) |
| $VAF(M_i, \mathcal{B}_s)$ | variant allele frequency of SNV $M_i$ in sample $\mathcal{B}_s$ (defined as the ratio $v_{is}/t_{is}$) |
| $\mathcal{F}(C_j, \mathcal{B}_s)$ | fractional copy number of allele A in sample $\mathcal{B}_s$ (obtained from the output of a tool for inferring tumor subclonal composition with respect to copy number states, see example in Figure 1d) |
| $\mathcal{M}(C_j)$ | set of all SNVs in CNA-consistent segment $C_j$ from the set $C - C'$ |
| $\phi_{vs}$ | cellular prevalence of subclone corresponding to node $v$ of tree $T$ in sample $\mathcal{B}_s$ |
| $\delta_{iv}$ | binary variable set to 1 if and only if SNV $M_i$ occurs for the first time at node $v$ |
| $\theta_{kv}$ | binary variable set to 1 if and only if copy number state $S_k$ occurs for the first time at node $v$ |
| $\mathcal{T}_{Av}$ | non-negative integer variable encoding the total number of copies of allele A at node $v$ |
| $\psi_{vk}$ | binary variable set to 1 if and only if copy number status at node $v$ is $S_k$ |
| $\mu_{iA}$ | binary variable set to 1 if and only if SNV $M_i$ occurs for the first time on allele A |
| $A_{iv}$ | non-negative integer variable encoding the number of copies of allele A that harbor SNV $M_i$ at node $v$ |
| $e_{Av}$ | binary variable defined for each non-root node $v$ and set to 1 if and only if $\mathcal{T}_{Av} = \mathcal{T}_{Ap(v)}$ |
| $l_{Av}$ | binary variable defined for each non-root node $v$ and set to 1 if and only if $\mathcal{T}_{Av} < \mathcal{T}_{Ap(v)}$ |
| $g_{Av}$ | binary variable defined for each non-root node $v$ and set to 1 if and only if $\mathcal{T}_{Av} > \mathcal{T}_{Ap(v)}$ |
| $I_{iv}$ | binary variable set to 1 if and only if there is at least one copy of allele A or B harboring $M_i$ at $v$ |
| $U_{Aiv}$ | binary variable set to 1 if and only if all copies of allele A at node $v$ harbor $M_i$ and $\mathcal{T}_{Av} > 0$ |

### C Additional details about simulated and real data analysis

#### C.1 Obtaining baseline tree on the set of SNVs from regions not impacted by CNAs

To obtain baseline tree of tumor progression on the set of SNVs from regions not impacted by CNAs, we first clustered such SNVs based on similarity in their read counts across samples [30], and then ran B-SCITE [16] on the set of the inferred clusters. B-SCITE was originally developed as a method for integration of single-cell and bulk sequencing data, but it can also build trees using either of the two data types alone. In particular, to infer a tree on bulk data alone it suffices to set weight parameter *w* (which weights contribution of single-cell data) to 0 (an artificial values for single-cell related data can be provided since they are required parts of the input, but they will not have an impact on tree inference provided the above setting of *w*). We ran B-SCITE for 100000 iterations and 3 repeats.

#### C.2 Details of running DETOPT, DeCiFer, CITUP and PhyloWGS Details of running DETOPT

Full details and instructions on running `DETOPT` are provided in the repository, which is available at the following link: https://github.com/algo-cancer/DETOPT. Note that in all instances presented in this work `DETOPT` was run using the weight parameter *ρ* set to 0.25 (see the ILP objective function provided in Section 2.4 for definition of this parameter).

##### Details of running DeCiFer

Two files, containing the information about SNVs (`input` `·tsv`) and tumor purity (`purity` `·txt`), in the format given in the DeCiFer input data specifications, were provided as the inputs to this tool. For all simulated instances, we provided the ground truth values of tumor purity. We ran DeCiFer for 200 restarts and, for the sake of reproducibility, we fixed the random seed to 42. Minimum and maximum number of clusters, as well as elbow criterion parameters were set experimentally in order to avoid both underclustering and overclustering. We observed that DeCiFer achieves the best performance when parameters `MINK` and `MAXK` were respectively set to 2 *·* (*h* + 1) and 2*·*`size of simulated tree`, and elbow criterion parameter to 0.008. The full command used to run DeCiFer is given below:

~~~
decifer input.tsv -p purity.txt -k MINK -K MAXK --elbow 0.008 --seed 42 -j 10 -r 200
~~~

##### Details of running CITUP

CITUP is a tree inference method that is typically used in combination with SNV clustering algorithms, which are run first in order to group SNVs based on similarity in their read counts across samples while accounting and correcting for the impact of CNAs. DeCiFer is one of the most recently developed clustering algorithms that works on multi-sample bulk sequencing data and provides descendant cell fraction values for each SNV in each sample, which can then serve as input to CITUP for building tree of tumor progression. As described in [30], descendant cell fractions are in general more suitable for tree inference than cellular prevalence values reported by some other clustering algorithms [27], especially in cases when SNVs are impacted by losses. After clustering variants using DeCiFer, we ran CITUP on the inferred set of clusters and providing CITUP with the ground truth tree topology. Since the number of clusters inferred by DeCiFer can differ from the size of the ground truth tree, we also ran B-SCITE (see Supplementary Section C.1) in order to infer tree of the same size as the number of clusters reported by DeCiFer. Since the first combination (i.e., running DeCiFer followed by CITUP with the ground truth tree provided) yielded results of higher accuracy only those results are shown in the paper.

##### Details of running PhyloWGS

Two files, containing the information about SNVs (`s` · `txt`) and CNAs (`c` · `txt`), in the format described in the PhyloWGS input data specifications, were provided as the inputs to this tool. Number of burn-in and true MCMC samples were respectively set to the default values, 1000 and 3500. Number of chains were set to 7. The full command used to run PhyloWGS is given below:

~~~
python multiEvolve.py --num-chains 7 --ssms s.txt --cnvs c.txt --burnin-samples 1000 --mcmc-samples 3500
~~~

As per recommendations in the PhyloWGS GitHub repository, we selected a tree with the highest LLH (log likelihood) to compute Ancestor-Descendant and Different-Lineage accuracy scores. We set the maximum run-time for PhyloWGS to 5 days (120 hours); when the time elapsed, any running jobs were cancelled and we considered these as instances where the tool failed to converge.

#### C.3 Details of real data analysis

Files containing additional details about the real data and all inputs and outputs of `DETOPT` can be found in the repository https://github.com/algo-cancer/DETOPT in folder `real data`. Specifically, for checking longitudinal consistency mentioned in the main text, please refer to the subfolder `longitudinal consistency`. Complete lists of assignments of variants to tree nodes can be found in the subfolder `variants placements`.

For inferring subclonal architecture with respect to copy number aberrations, we used the latest version of HATCHet available at https://github.com/raphael-group/hatchet. All outputs of HATCHet are available in the subfolder `hatchet output` of the `real data` folder in `DETOPT`’s repository.

## D Supplementary Figures

**Supplementary Figure 1:**
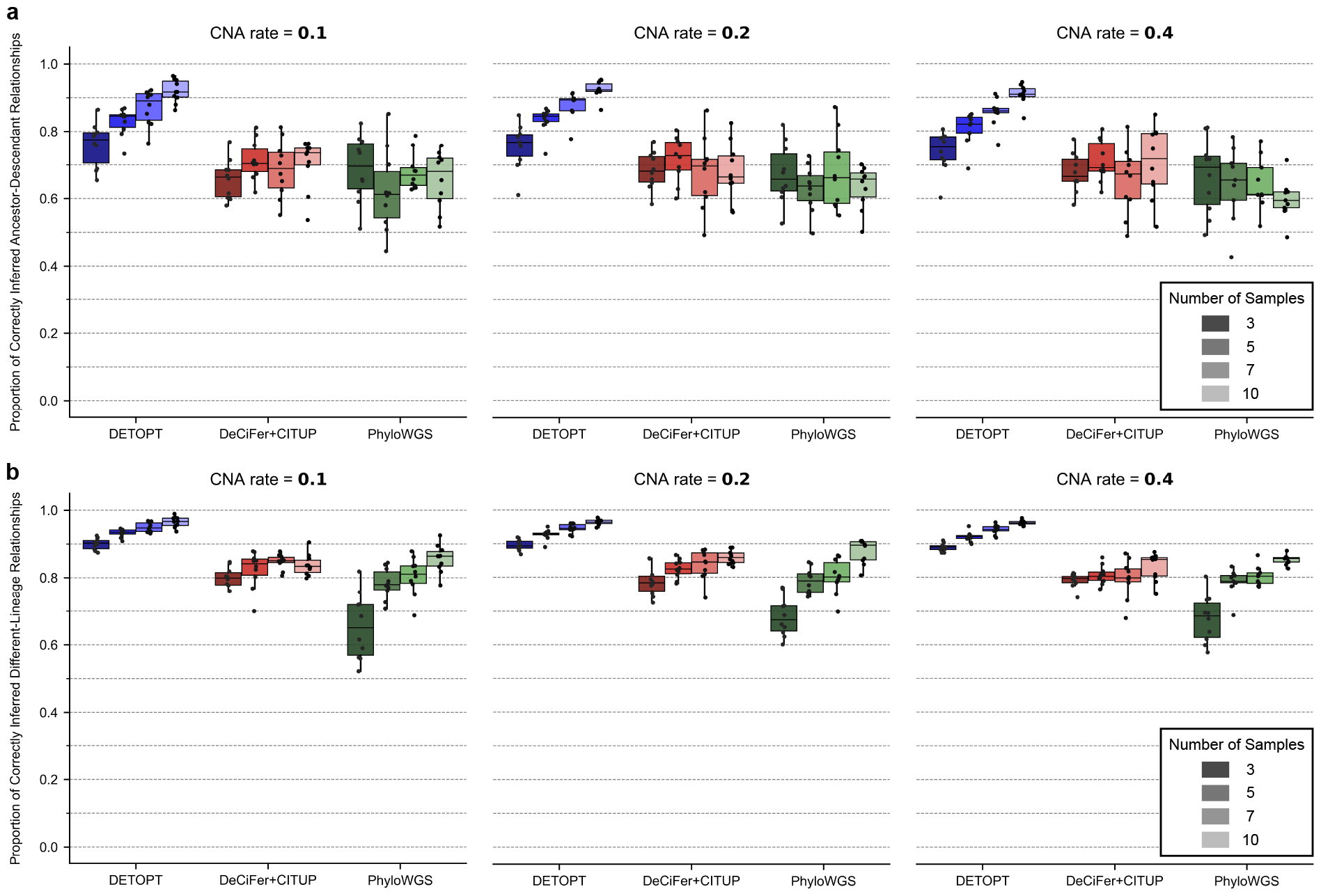
Comparison of `DETOPT`, DeCiFer+CITUP, and PhyloWGS on simulated data with larger tree sizes (i.e., number of subclones). All parameters are same as in Figure 2 in the main text, except that the tree size was set to 20. For each parameter combination, 10 different trees were simulated. Plotted are the **a**. Ancestor-Descendant accuracy (see the main text for definition). **b**. Different-Lineage accuracy (see the main text for definition). PhyloWGS failed to converge in 3 (out of 120) instances within the time limit of 120 hours and no results for this method are shown for those 3 instances.

**Supplementary Figure 2:**
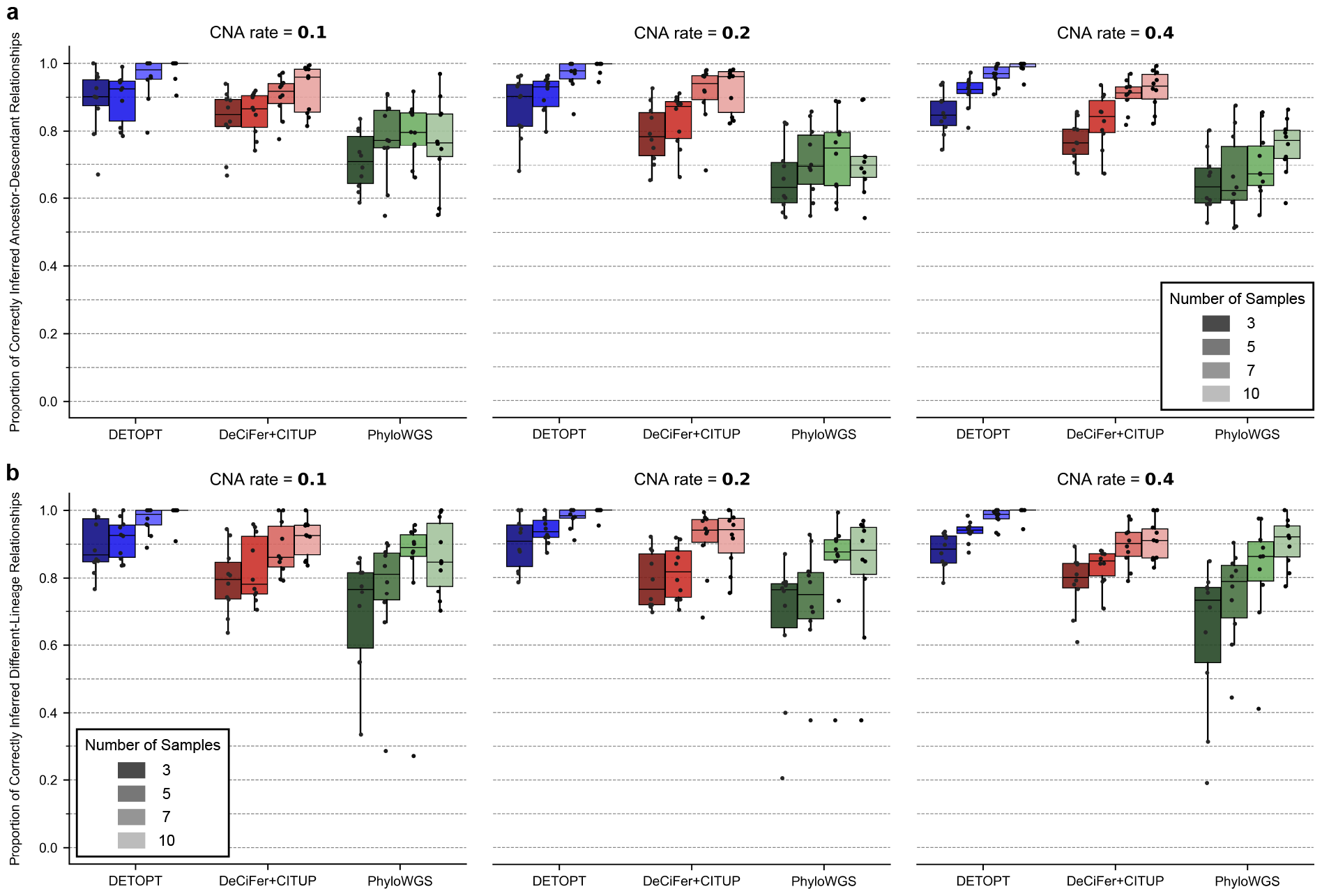
Comparison of `DETOPT`, DeCiFer+CITUP, and PhyloWGS on simulated data with lower number of SNVs. In all simulated instances shown, the total number of SNVs was set to 100 with varied proportions of them being impacted by a CNA (0.1, 0.2, and 0.4), tree size was set to 10, and sequencing coverage to 100*×*. For each parameter combination, 10 different trees were simulated. Plotted are the **a**. Ancestor-Descendant accuracy (see the main text for definition). **b**. Different-Lineage accuracy (see the main text for definition).

**Supplementary Figure 3:**
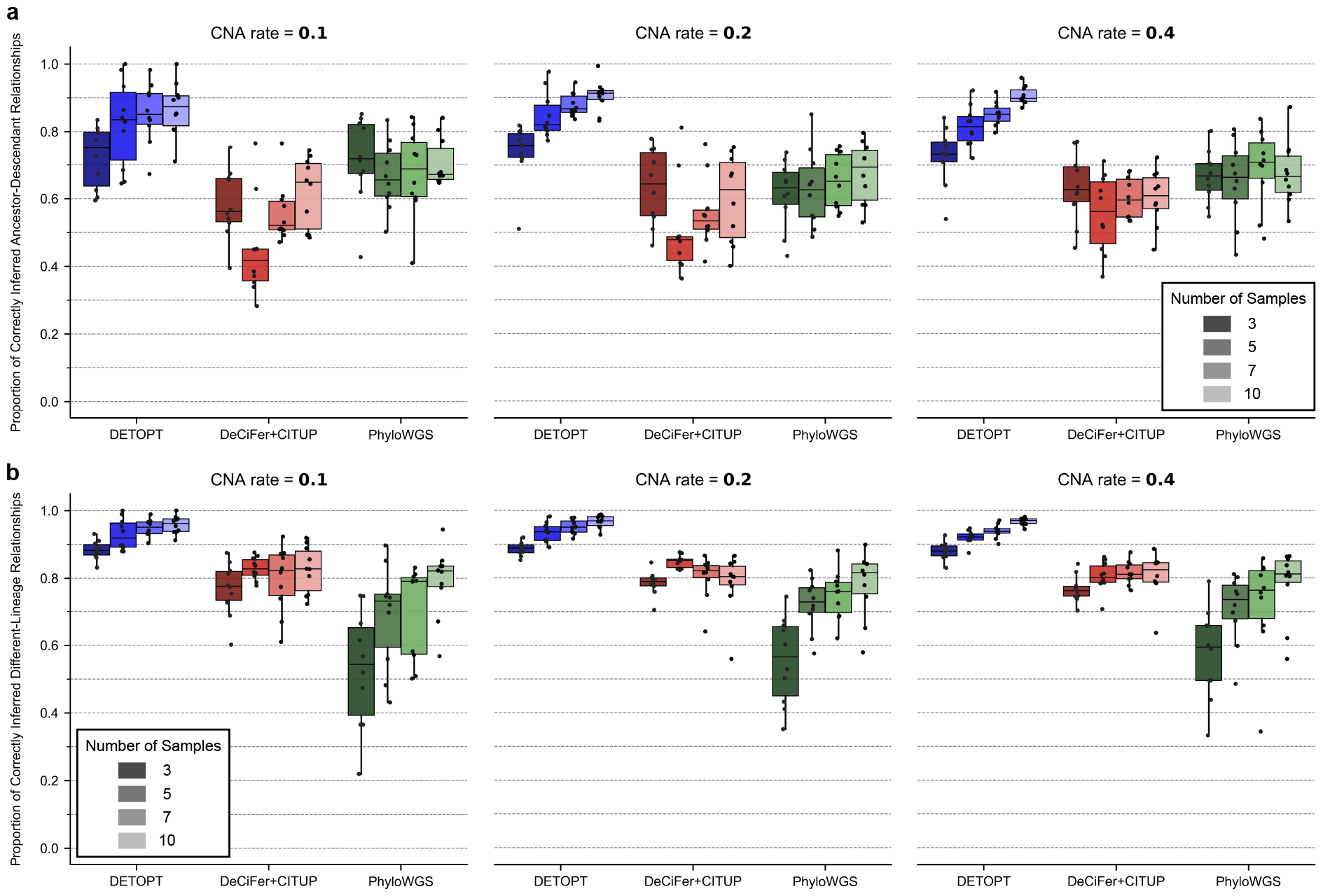
Comparison of `DETOPT`, DeCiFer+CITUP, and PhyloWGS on simulated data with larger tree sizes. All parameters are the same as in Supplementary Figure 2, except that the simulated trees were of size 20. For each tool, plotted are the **a**. Ancestor-Descendant accuracy (see the main text for definition). **b**. Different-Lineage accuracy (see the main text for definition).

**Supplementary Figure 4:**
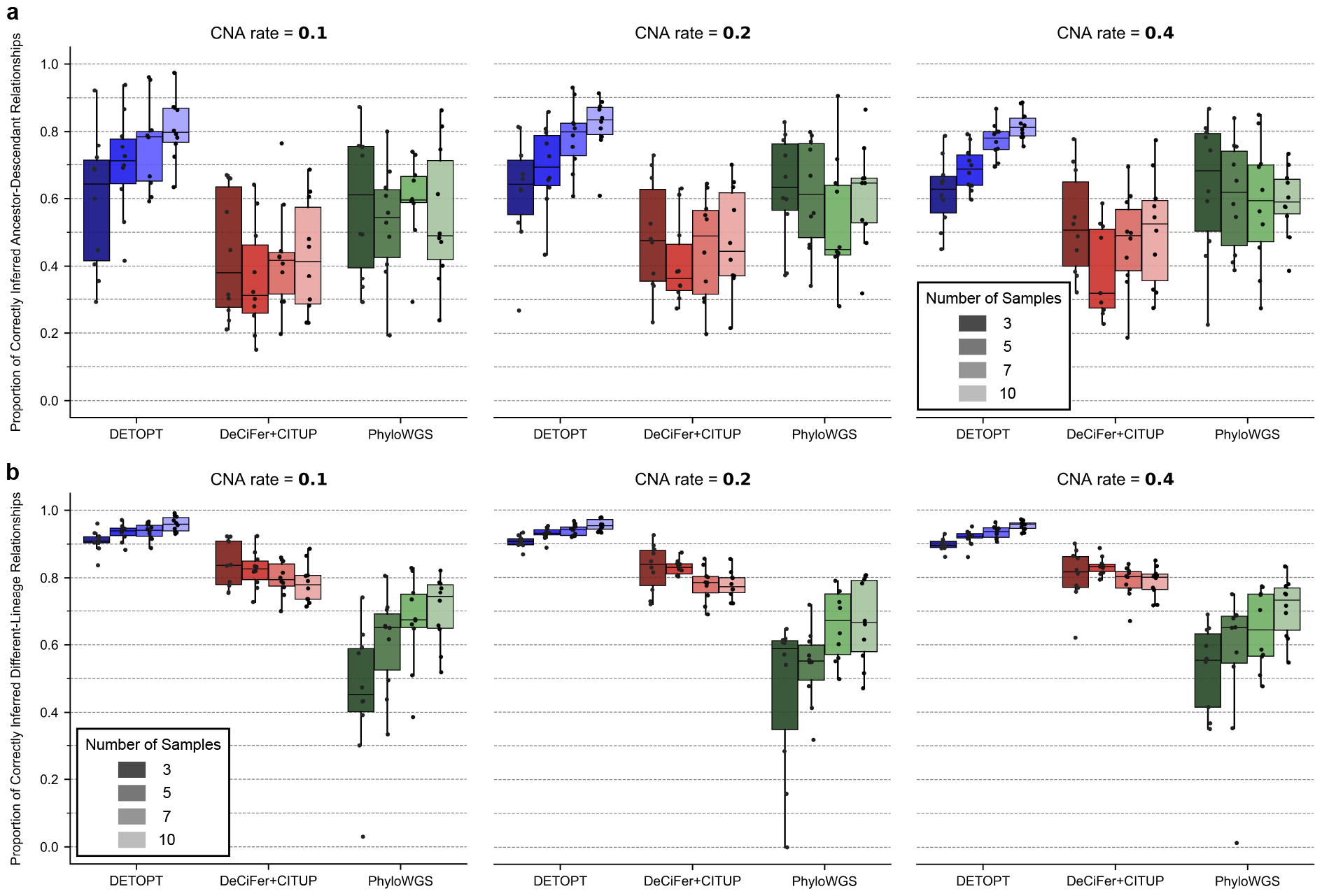
Comparison of `DETOPT`, DeCiFer+CITUP, and PhyloWGS on simulated data with very large tree sizes. All parameters are the same as in Supplementary Figure 3, except that the simulated trees were of size 30. For each tool, plotted are the **a**. Ancestor-Descendant accuracy (see the main text for definition). **b**. Different-Lineage accuracy (see the main text for definition).

**Supplementary Figure 5:**
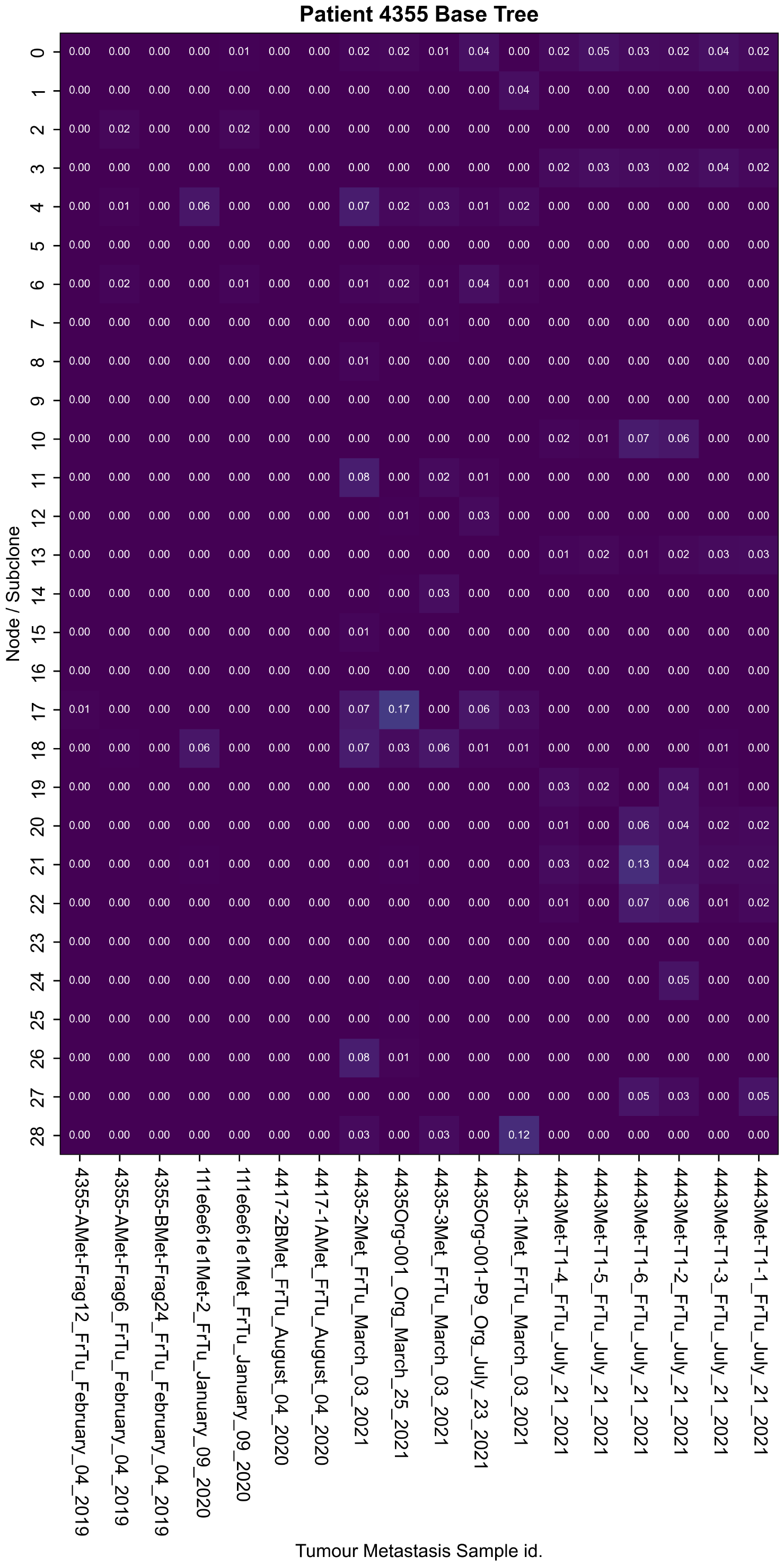
Heatmap showing absolute values of differences between the inferred and observed cellular prevalence values of mutational clusters used in building baseline tree for mBrCa patient presented in the main text. Observed cellular prevalence of a cluster is computed by averaging cellular prevalence values of SNVs belonging to the cluster (estimated as 2 · *V AF*). Rows correspond to samples and columns to clusters.

## Footnotes

1 Note that the summary of all symbols and variables used in this section is provided in Supplementary Table 1.

2 This assumption is plausible in this context as it is unlikely that both homologous copies of a chromosomes independently acquire the very same SNV; even if such an event could occur, it would be rare and statistically insignificant and would not have a major impact on tree reconstruction.

3 Note that here we assume that | *𝒮* (*C*_*j*_) | ≤ |*V* (*T*)|, which is plausible assumption as tree size is usually significantly larger than the number of distinct copy number states of a given segment.

4 Note that these and several other constraints below are quadratic in the presented form, but they can be linearized using standard linearization techniques for product of two non-negative integer variables.

